# *Tulp3* quantitative alleles titrate requirements for viability, brain development, and kidney homeostasis but do not suppress *Zfp423* mutations in mice

**DOI:** 10.1101/2025.04.25.650726

**Authors:** Corinne A. McCoy, Dorothy Concepcion, Mark G. Mezody, Raquel Z. Lara, Ojas Deshpande, Catherine Liang, Renee Long, Bruce A. Hamilton

## Abstract

Tubby-like protein 3 (TULP3) regulates receptor trafficking in primary cilia and antagonizes SHH signaling. *Tulp3* knockout mice are embryonic lethal with developmental abnormalities in multiple organs, while tissue-specific knockouts and viable missense alleles cause polycystic kidney disease. Human patients with *TULP3* mutations present with variable, but often multi-organ fibrotic disease. We previously showed that mouse and human *Tulp3* expression is negatively regulated by ZNF423, which is required for SHH sensitivity in some progenitor cell models. The level of TULP3 function required to prevent mutant phenotypes has not been known. Here we report a *Tulp3* quantitative allelic series, designed by targeting the polypyrimidine tract 5’ to the splice acceptor of a critical exon, that shows distinct dose-response effects on viability, brain overgrowth, weight gain, and cystic kidney disease. We find limited evidence for genetic interaction with *Zfp423* null or hypomorphic mutations. Together, these results establish an approach to developing quantitative allelic series by exon exclusion, rank-order dose-sensitivity of *Tulp3* phenotypes, and model thresholds for TULP3 function to prevent severe outcomes.

**Author Summary:** TULP3 protein plays critical roles in regulating receptor trafficking and signaling in the primary cilium. Mutations in the *TULP3* gene can cause severe, multi-organ disorders in both mice and humans, yet the amount of *TULP3* activity needed to avoid these outcomes has been unclear. In this study, we used precise genome editing in mice to create a set of new *Tulp3* gene variants that reduce TULP3 expression to varying degrees. This allowed us to test how much TULP3 is required for survival, normal brain and kidney development, and weight regulation. We found that as little as 5% of normal *TULP3* levels is enough to avoid lethal birth defects, but still leads to obesity, mild brain overgrowth, and progressive kidney cysts preceded by reductions in cilium frequency and length in situ. The severity of these effects was related to TULP3 protein levels, highlighting a dose-dependent response. We also investigated whether reducing TULP3 levels would suppress brain abnormalities in *Zfp423* mutant mice, based on prior evidence of a genetic interaction, but did not find evidence to support this effect. Our work provides a framework for understanding how varying levels of *TULP3* affect various organ systems and offers a general strategy for creating quantitative genetic models of human disease.

## INTRODUCTION

TUB-like protein 3 (TULP3), like TUB, is a phosphatidylinositol-(4,5)-bisphosphate (PIP2) binding protein that is dynamically localized to different cell compartments, including plasma membrane, nucleus, and primary cilia, during G protein coupled receptor signaling [1, 2]. TUB and TULP3 are broadly expressed, close homologs [3] that promote trafficking of membrane proteins into primary cilia through interactions with phosphatidylinositol-(4,5)-bisphosphate (PIP2), the intraflagellar transport complex A (IFT-A), and cargoes–notably G protein-coupled receptors and polycystins [4–10]. TULP3 antagonizes SHH signaling through its trafficking activity [2, 4, 6, 11–13], with consequences for the development and function of several organ systems. We previously showed that *Tulp3* is a target of *Zfp423* in both mouse and human cells [14] and set out to test their genetic interaction *in situ*.

Mouse genetics identified functional requirements for *Tulp3* across several organs and cell types. *Tulp3* mutants from either ES-cell targeting or chemical mutagenesis showed embryonic lethality with exencephaly, spina bifida, preaxial polydactyly, splayed vertebrae, and increased SHH activity [2, 12, 15]. A subsequent viable missense allele, *Tulp3^K407I^* (a.k.a. *Tulp3^m1Kflj^*), and conditional deletion in either collecting ducts (*Tulp3^tm1a(EUCOMM)Hmgu^* with *Hoxb7-*Cre) or nephrons (with *Ksp*-Cre) both showed polycystic kidneys in embryos or young mice, while simultaneous deletions of *Tulp3* and the polycystin gene *Pkd1* suppressed the renal cysts seen in either single gene deletion [16, 17]. Despite this progress, the amount of *Tulp3* function required for viability and the dose-response for postnatal phenotypes have remained open questions and direct comparisons across alleles in different labs are complicated by use of different genetic backgrounds for each of these mutations.

Subsequent human studies identified biallelic damaging *TULP3* variants in patients with hepatic fibrosis, fibrocystic kidneys, cardiomyopathy, and compromised DNA damage response [18], which is a shared feature among several ciliopathy gene disorders [19–23]. Variants included frameshift, nonsense, splice variant, and amino acid substitutions in the highly conserved TUB domain. An additional patient reported later in ClinVar had a splice donor mutation after exon 7. Onset and progression were variable among affected individuals in the documented cohort, but liver and kidney damage preceded other organ involvement, with non-obstructive hypertrophic cardiomyopathy reported only in a minority of patients and only in the sixth decade or later.

Typical for rare genetic disorders, how much of the variation in disease presentation is due to allelic differences and how much to other causes–and how much functional recovery would be required for benefit–are not known. Recent evidence has also implicated TULP3 in the effects of caloric restriction on aging [24].

Given high levels of genetic heterogeneity (both locus and allelic) and variable penetrance of disease phenotypes in human ciliopathies as well as differing genetic backgrounds and individualized phenotyping protocols in animal models, relatively few studies have isolated allele-specific effects from potential effects of genetic background and non-genetic factors.

Fewer have done so with quantitative alleles that titrate levels of genetic activity at the subject locus. Here we develop a *Tulp3* quantitative allelic series in mice by editing the polypyrimidine tract adjacent to a frame-shifting exon (exon 7) to modulate exon inclusion and expression level, as well as qualitative variants lacking a conserved Tub-N domain (exon 4). We titrate clinically relevant phenotypes across this series and test prior hypotheses for genetic interaction between *Tulp3* and *Zfp423* mutations in mice. Our results establish a generalizable approach and show that minimal (≤5%) of TULP3 expression level is required for viability, that modest increments of expression improve clinically-related phenotypes, and that for at least one exon, loss of conserved protein segments has a comparatively modest impact on genetic function.

## RESULTS

### Quantitative allelic series by targeting inclusion of a frame-shifting exon

We targeted two sites in *Tulp3* based on its exon structure and protein coding domains, using a high-fidelity Cas9 and a pool of donor oligonucleotides to template repair on a coisogenic FVB/NJ genetic background (Figure 1A). We generated two mutations at exon 4, which encodes a conserved protein domain, but whose skipping would maintain the open reading frame in the resulting mRNA (Figure 1B). Both mutations should abolish exon 4 inclusion in *Tulp3* mRNA but have residual genetic function unless the conserved Tub-N domain is essential to all TULP3 activities. We targeted the polypyrimidine tract 5’ to *Tulp3* exon 7 splice acceptor to decrease the exon inclusion rate in processed mRNA. Exon 7 was selected based on its exclusion resulting in a reading frame shift, such that the rate of exon 7 inclusion should predict the degree of genetic function of an edited allele. We recovered seven distinct alleles from a single injection job, including all four templated by the oligonucleotide pool and three insertion/deletion (indel) variants, of which two (Δ5, Δ3) were predicted to abolish exon 7 inclusion and function as null mutations (Figure 1C). Oligonucleotide donors for editing and assay primers are given in S1 and S2 Tables.

**Figure 1.**
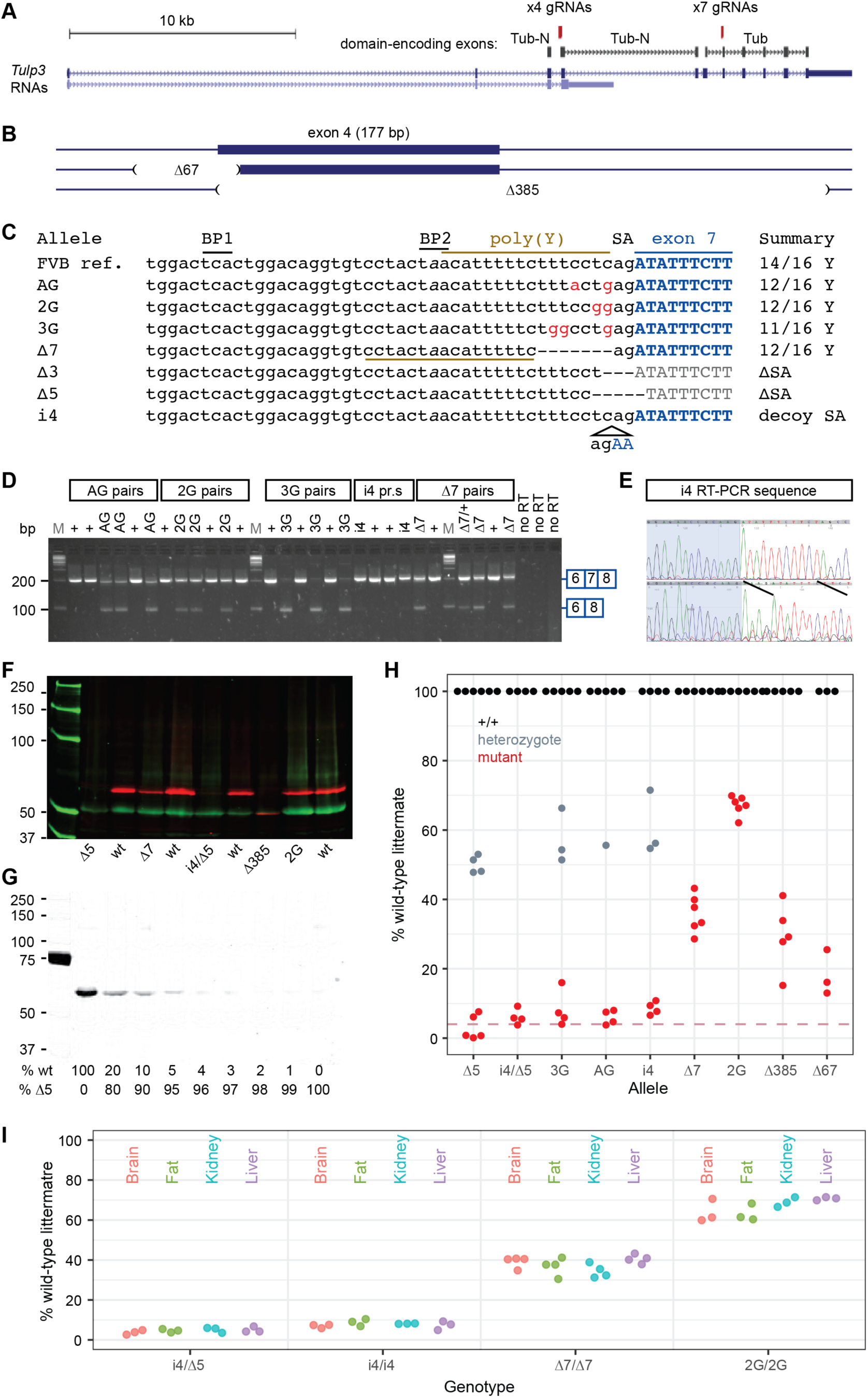
Allelic series targeting polypyrimidine tracts. (**A**) Genomic organization of *Tulp3* showing targeted exons relative to encoded domains. (**B**) Deletion alleles relative to exon 4. (**C**) Exon 7 alleles relative to FVB/NJ reference sequence. BP, splicing branch points; SA, splice acceptor; poly(Y), polypyrimidine tract. (**D**) RT-PCR shows allele-dependent levels of exon skipping induced by mutations in the polypyrimidine tract 5’ to exon 7. (**E**) Sanger sequence of a control (top) and *i4* homozygote (bottom) shows a majority of steady-state RNA spliced into the mutant-specific decoy splice acceptor. (**F**) Two-color Western blot detects TULP3 (red, upper) and loading control (green, lower) from E14.5 heads relative to size marker (kD). (**G**) Null (Δ5/Δ5) protein extracts with wild-type spike-in estimates detection limit at ∼4% of control. (**H**) Band intensities relative to littermate controls define a quantitative allelic series based on reduced TULP3 protein expression. Dashed line, 4%. (**I**) Band intensities relative to littermate controls for show do not find nominally significant differences across four tissues at P29 (p-values 0.11 to 0.56 for each genotype, one-factor ANOVA).

RT-PCR assays show that the exon 7 mutations result in reduced exon 7 utilization. Exon inclusion assays showed inclusion rates that quantitatively distinguished alleles (Figure 1D). The non-templated 4-bp insertion allele (i4) predicted a strong alternative splice acceptor that would produce a frameshift without inducing exon skipping. DNA sequencing of the RT-PCR product from an *i4* homozygote of steady-state *Tulp3* RNA is spliced into this upstream decoy site rather than the in-frame splice acceptor (Figure 1E).

Western blots confirmed and quantified reductions in TULP3 protein expression across the allelic series. Extracts from E14.5 embryo littermate pairs showed reduced expression of otherwise wild-type TULP3 protein from each of the exon 7 alleles, with a reduced steady-state amount of a smaller TULP3 protein from the exon 4-skipping alleles (Figure 1F). Spike-in controls showed a detection limit at ∼4% of wild-type levels under our conditions (Figure 1G). The very low levels achieved by splicing variants and null heterozygote expression at 50% suggest little to no compensatory feedback regulation for TULP3 levels. The lowest expression genotype, i4/Δ5 compound heterozygotes, expressed ≤5% of wild-type littermate TULP3 levels (Figure 1H and S3_Table). For viable genotypes, expression level did not vary significantly among four tissues relevant to *Tulp3* phenotypes at postnatal day 29 (Figure 1I and S4_Table). These results demonstrate simple construction of a quantitative allelic series that can be used for calibrating genetic function relative to *TULP3* phenotypes or potential interventions, using an approach that should be generalizable to other genes and across organ systems.

### Minimal TULP3 is required for survival to adulthood

Having a quantitative allelic series on a coisogenic background allowed us to estimate the amount of *Tulp3* genetic function required to escape lethality and other major phenotypes. The Δ5 splice acceptor mutation behaved as a genetic null. Homozygous Δ5 embryos produced no detectable TULP3 protein and showed brain overgrowth resulting in exencephaly and preaxial polydactyly resulting in supernumerary digits by E14.5 (Figure 2A), consistent with previous knockout models on other strain backgrounds [2, 11, 12, 15]. Postnatally, we found no surviving homozygotes among ∼200 progeny from null-allele heterozygote crosses (Figure 2B). By contrast, hypomorphic genotypes, including the i4/Δ5 allelic combination that expressed only ∼5% of normal TULP3 levels showed no significant evidence for reduced viability in moderately large crosses. A nominal effect of the least severe allele, 2G, did not survive correction for family-wise error rate and included the caveat that allele-specific PCR assays for 2G, 3G, and AG were tuned to avoid homozygote false-positives and therefore likely under-count true homozygotes. Pooling hypomorphic allele data from all length-specific assays (i4/Δ5, i4, Δ7, and the two exon 4 deletions) was not significant despite large aggregate sample (p=0.17, Chi-squared test). Littermate pairs selected for age-dependent phenotypes similarly failed to support a significant effect of hypomorphic genotypes on later survival (likelihood ratio test p = 0.9 for i4/Δ5, n = 29 littermate pairs to 150 days; p = 0.5 for combined pool of i4/Δ5, i4/i4, 3G/3G, and AG/AG strong alleles, n = 84 littermate pairs; p = 0.5 for exon 4 deletion, n = 61 littermate pairs; p = 0.2 for 248 littermate pairs across all variants). These results suggest that for severe *Tulp3* phenotypes, even small increments of improved function might be beneficial, with only ∼5% of wild-type levels required for gestation and long-term survival.

**Figure 2.**
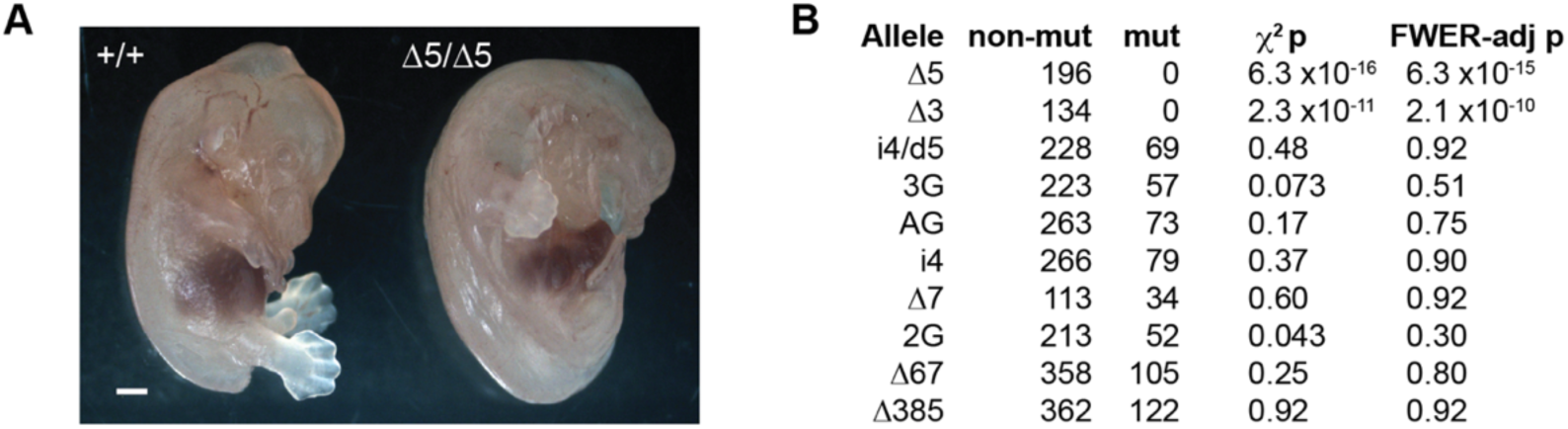
Minimal TULP3 expression is sufficient for viability and survival to adulthood. (**A**) Exencephaly and polydactyly are evident by E14.5 in Δ5 homozygotes. Bar, 1 mm. (**B**) Genotype ratios show complete lethality of null animals, but no significant evidence for reduced viability among even the strongest hypomorphic genotypes with TULP3 expression near the limit of detection in Western blots. Nominal p-values from the 1-df chi-squared test were adjusted for family-wise error rate (FWER-adj p) using Hommel’s method [25].

### Hypomorphic *Tulp3* genotypes gain excess weight

From littermate pairs left during several months of the COVID-19 pandemic, *Tulp3* hypomorphic mutants were visibly obese relative to co-housed same-sex littermates (Figure 3A and S5_Table). As a more quantitative test and to determine the time course for onset, we followed co-housed, same-sex littermate mutant-control pairs with periodic weight measurements in a subsequent cohort from weaning to >6 months. For the lowest TULP3-expressing genotype (i4/Δ5), a clear trend showed increasing weight gain over time by mutants relative to their littermate controls for both sexes (Figure 3B). Results from other mutants showed similar though less pronounced trends, roughly proportional to expression level and loss of exon 4 (Figure 3C). As some littermate pairs were censored at later times due to age-related health issues independent of genotype, we examined pairs weighed at 100-132 days postnatal (Figure 3D).

**Figure 3.**
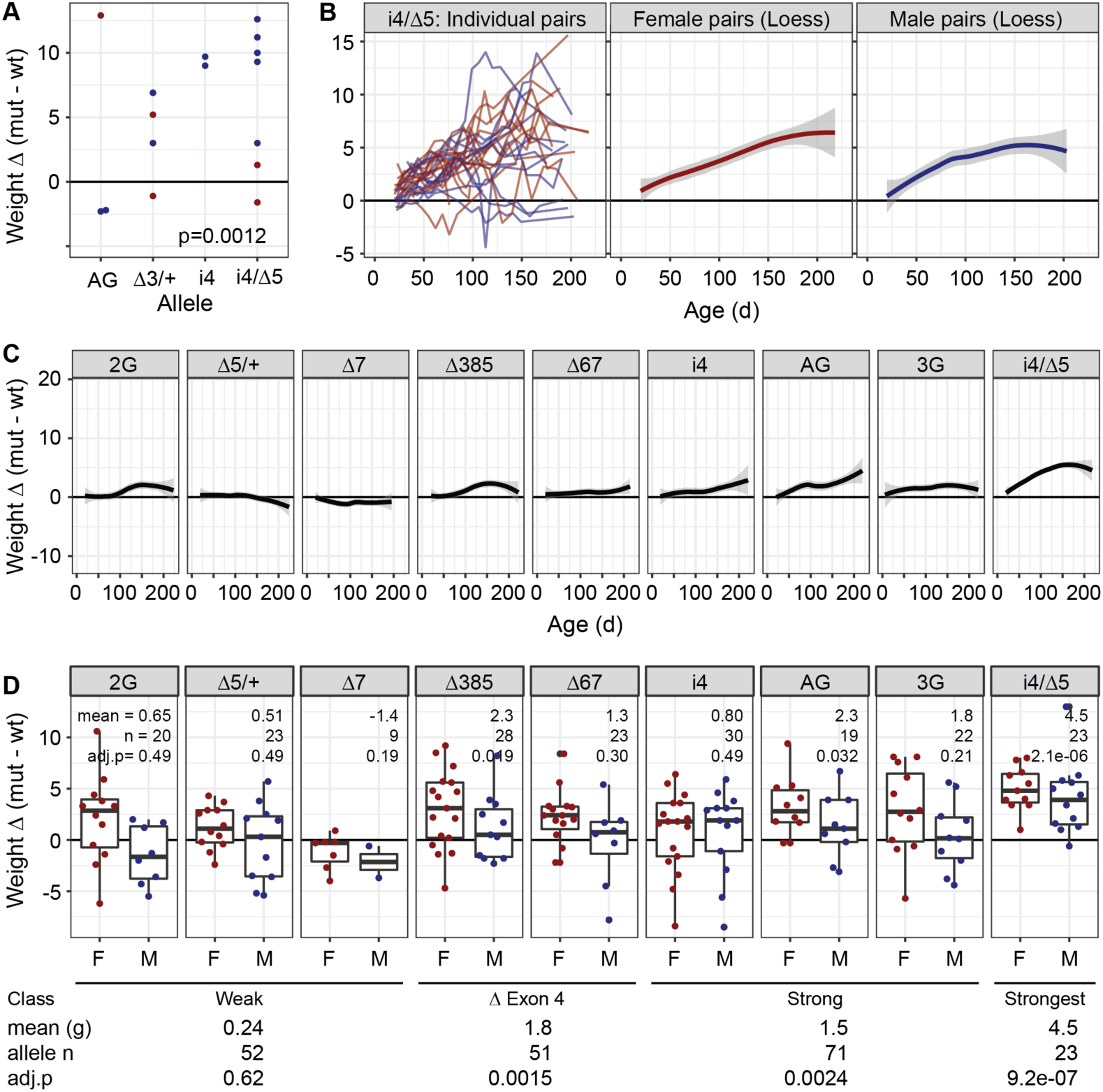
Allele-dependent weight gain in *Tulp3* hypomorphs. (**A**) Incidental observation of co-housed sex-matched littermates showed obesity in aged *Tulp3* mutants. Each point is the difference (mutant-wildtype) in grams for a littermate pair. Mean age 220 days. (**B**) Time course of weight difference in individual i4/Δ5 littermate pairs and loess regression plots of i4/Δ5 pairs for each sex. (**C**) Loess plots for each allele, both sexes. (**D**) Summary boxplots with points for distinct individual pairs (first measure after postnatal day 100) for each *Tulp3* allele. Mean difference in grams, sample size, and FWER-corrected p-values from paired sample t-tests are given for each allele (above) and expression class (below).

While modest sample sizes and corrections for family-wise error rate limited power for individual genotypes, aggregating variants whose effect on TULP3 expression was relatively weak (2G, Δ5/+, and Δ7), weak but included removal of exon 4 (Δ67, Δ385), or relatively strong (i4, AG, 3G) showed a rough correlation between weight gain and TULP3 expression loss. We saw significant effects of all but the weak alleles, with exon 4 alleles roughly comparable to strong alleles, and with the i4/Δ5 combination having a stronger effect than all others. These data show a quantitative effect of TULP3 expression level on weight homeostasis at levels below 30% of wild-type.

### Strongest reduction in *Tulp3* produces increases in brain measures

*Tulp3* knockout mice, including Δ5 allele homozygote reported here, display multisystem disorders including exencephaly secondary to brain overgrowth (Figure 4 and S6_Table). To determine whether viable *Tulp3*-mutant mice display related changes to gross brain morphology, we adopted a pipeline of 8 simple measures (vermis width, cerebellar hemisphere anterior-posterior distance, and maximum brain width from dorsal photographs of whole brain; vermis sagittal area, cortical thickness, corpus callosum thickness, and anterior commissure thickness from block face preparation photographs) that we previously used for analysis of *Zfp423* allelic variants [26]. We found that i4/Δ5 compound heterozygotes, the most severe viable genotype with respect to TULP3 protein level, had increased vermis sagittal area and cerebellar hemisphere anterior-posterior distance (4.5-6.0%) after correction for multiple outcome measures, with nominal support for increased brain width (2.6%), relative to same-sex littermate controls (Figure 4). Pooled data from the next most severe genotypes (i4/i4, AG/AG, and 3G/3G) showed nominal increases in vermis area and brain width (1.3-2.1%) that did not survive correction for multiple tests. These data show that even viable reduction in TULP3 can result in modest brain overgrowth phenotypes and reinforce the conclusion that even modest increases in *Tulp3* function can significantly impact phenotypic outcomes.

**Figure 4.**
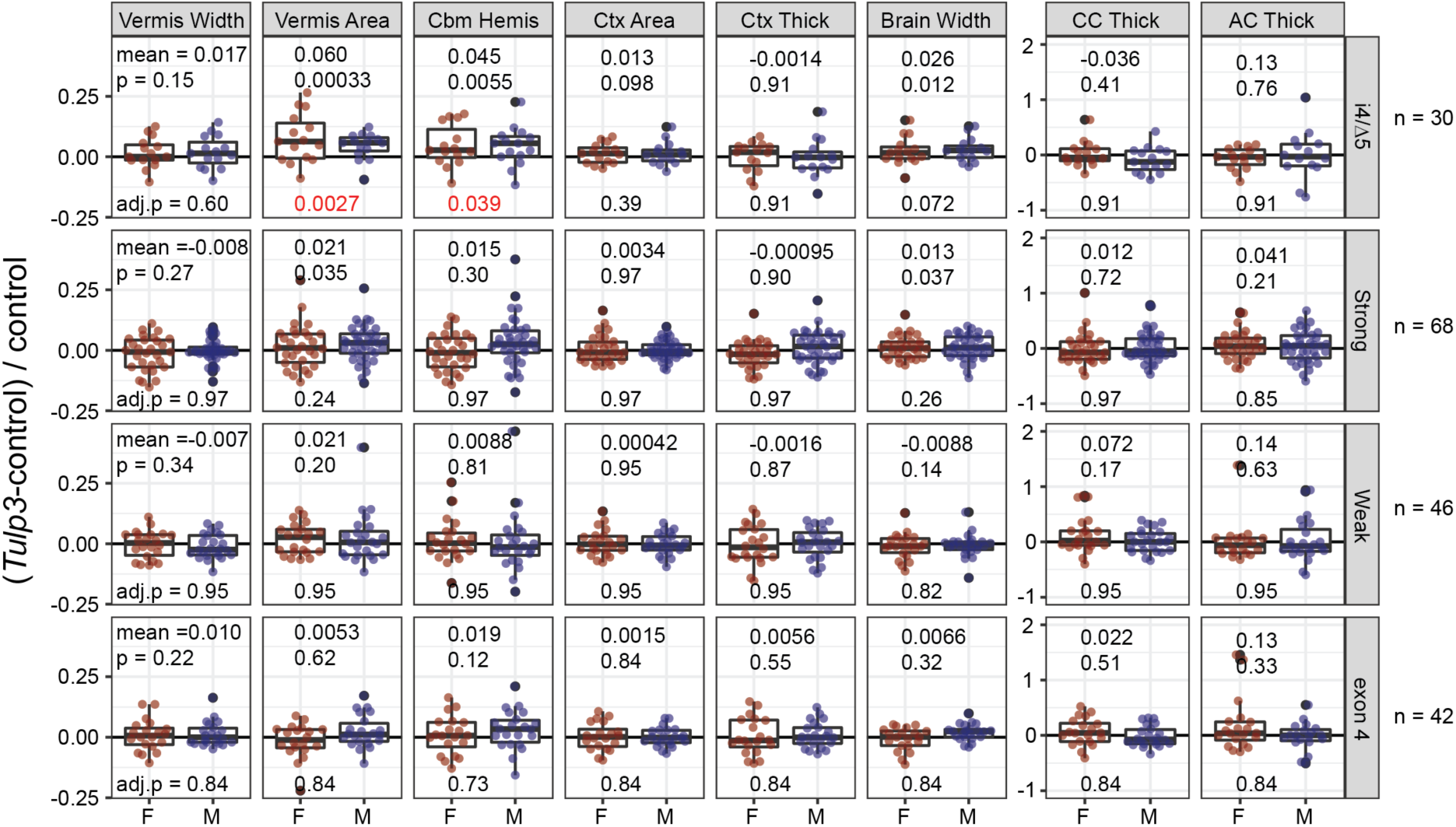
The strongest viable *Tulp3* variants influence brain morphology. (**A**) Compound heterozygotes i4/Δ5 average ∼5-6% larger cerebellar vermis sagittal areas and anterior-posterior dimension of the cerebellar hemispheres. The widest extent of the brain was also nominally larger, but did not survive correction for multiple phenotypes (top row). Other strong hypomorphic genotypes (i4/i4, 3G/3G, AG/AG; combined to improve power across a similar expression level) showed nominal support for a lesser increase in vermis area and brain width that did not survive correction (second row). Neither the weaker exon 7 alleles (2G and Δ7) nor the exon 4 deletions (Δ67, Δ385) returned even nominally significant results for any measure. Mean, nominal p-value (paired sample t or Wilcoxon signed rank test), and adjusted p-value after FWER correction across 8 phenotype measures are shown. Abbreviations: Cbm Hemis, anterior-posterior dimension of the cerebellar hemispheres; Ctx Area, cortical area in dorsal surface view; Ctx Thick, average cortical thickness at 15°, 30°, and 45° from midline in coronal block face view; CC and AC Thick, thickness of the corpus callosum and anterior commissure, respectively, at the midline.

### Progressive kidney cystogenesis responds to small increments of TULP3 expression

*Tulp3* tissue-specific knockouts and a viable missense variant had cystic kidney disease, but neither the time course of disease nor the quantitative level of gene function to reduce or ameliorate primary endpoints have been known. We quantified a cystic index for sectioned kidneys from same-sex littermate pairs across a range of ages for each *Tulp3* allele using CystAnalyzer [27] (Figure 5 and S7_Table). The presence and size of cysts in aged animals visibly varied across alleles (Figure 5A). Cysts were evident for each of the strong exon 7 alleles but less frequent or not evident for weaker alleles that expressed ≥30% of control levels of TULP3 (Figure 1G). The two exon 4 deletion/skipping alleles combined showed a reproducibly low level of cysts, consistent with the relatively low steady-state protein level we observed, but not consistent with exon 4 encoding an essential function. The extent of cystogenesis for each genotype inversely correlated with expression level and increased with age (Figure 5B). In a two-factor ANOVA model, genotype (p = 7.5 x 10⁻⁷) and age (p = 0.043) were significant, with significant pairwise differences between i4/Δ5 and all others (p < 1 x 10^-2^), and between i4 and 3G (p = 0.034) or weaker alleles, but i4 and AG were not statistically different (p > 0.99). Cystic index was both more sensitive and better correlated to TULP3 expression than simpler measures of kidney weight relative to body weight or kidney cross-sectional area (Supplemental S1_Figrue A-B). These data support a continuous relationship between kidney health and TULP3 expression below 50%, with particularly steep dose-response effects below ∼10% of control levels.

**Figure 5.**
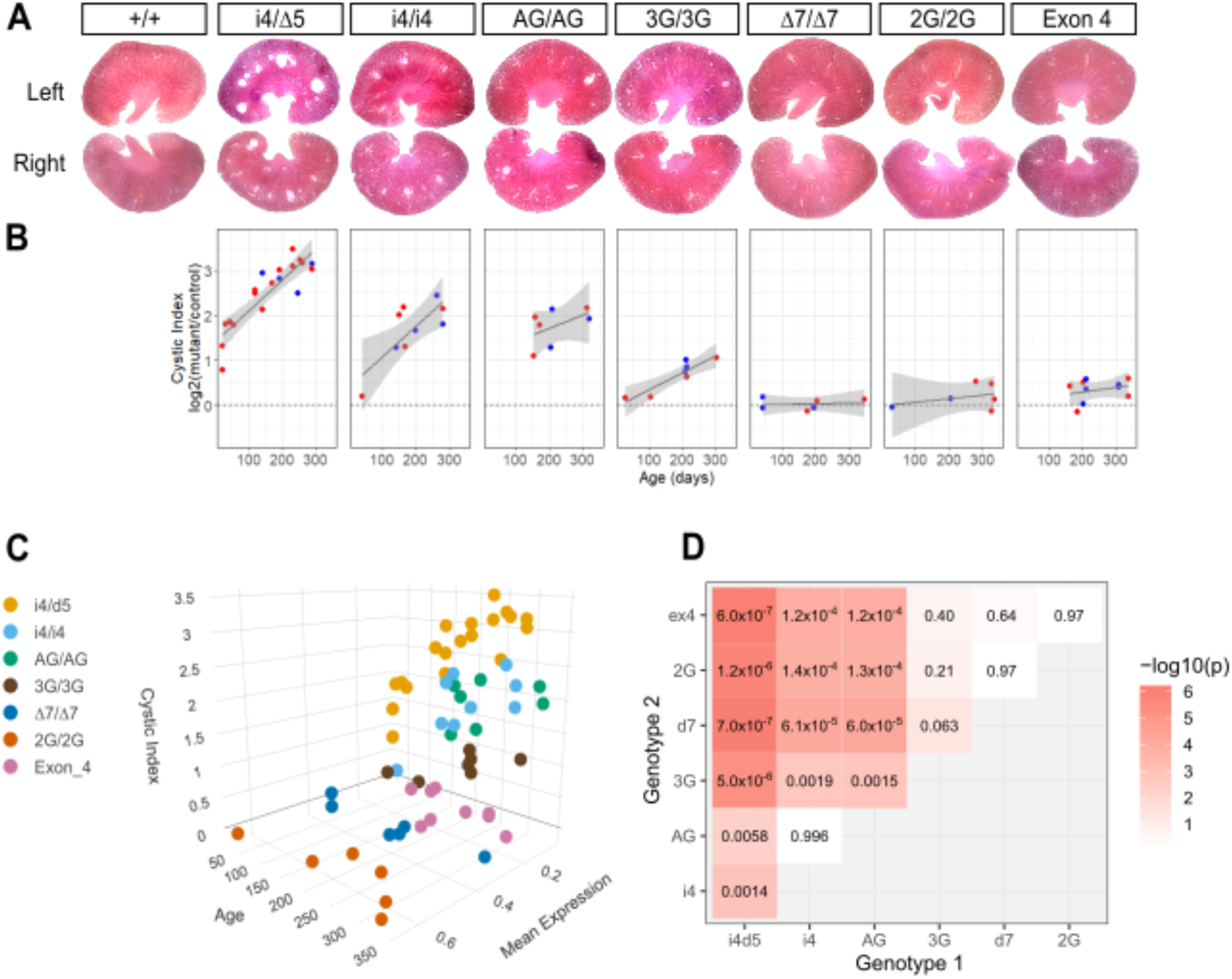
Allelic series shows dose-response in kidney cystogenesis. (**A**) Kidney sections from aged adult mice show allele-typical degrees of cystogenesis. (**B**) Cystic index for *Tulp3* variants showed allele- and age-dependence relative to same-sex control littermates. A generalized linear fit and 95% confidence interval are shown. (**C**) Three-dimensional visualization illustrates a sharp shift in cystic index across TULP3 expression levels. (**D**) Heatmap with printed values for TukeyHSD post-hoc test for difference between genotypes after two-factor ANOVA for cystic index as a function of genotype and age.

Despite evident tissue damage, veterinary blood chemistries from aged littermate pairs (minimum 119 days, mean 234) did not detect differences in standard measures across 16 sample pairs from strong hypomorphic alleles (Supplemental Figure S1_Figure and S8_Table).

This suggests that histological measures may be more sensitive than standard blood chemistries for modeling ciliopathy defects in mice.

### Reduced cilium length and frequency precede cystic kidneys

We next investigated whether reduced *Tulp3* expression or exon 4 deletion affected primary cilium structure and function in kidneys prior to cystogenesis, focusing on the most severe viable loss of expression (i4/Δ5) and exon 4-deleted (Δ67) models compared with their sex-matched littermates. We measured cilia length, frequency, and localization of TULP3 and GPR161 in kidney tissue from these mice at a pre-cystic stage. We measured length of 276 individual cilia based on immunofluorescence for ARL13B (Figure 6A, B and S9_Table). Mean cilium length was ∼15% shorter in i4/Δ5 compound heterozygotes than in control littermates across all four littermate pairs (p = 6.8 x 10^-5^, paired t-test) but not significantly different between Δ67 homozygotes and controls (p = 0.98). We also measured ciliation frequency for 500-600 nuclei per sample (Figure 6C and S10_Table). For i4/Δ5, ciliation frequency was significant decreased (OR = 0.81, p = 3.3 × 10^-6^, Fisher’s exact test), while for Δ67 mutants showed a modest but statistically borderline decrease (OR = 0.92, p = 0.059). These results suggested impaired ciliogenesis, with a stronger effect in i4/Δ5.

**Figure 6.**
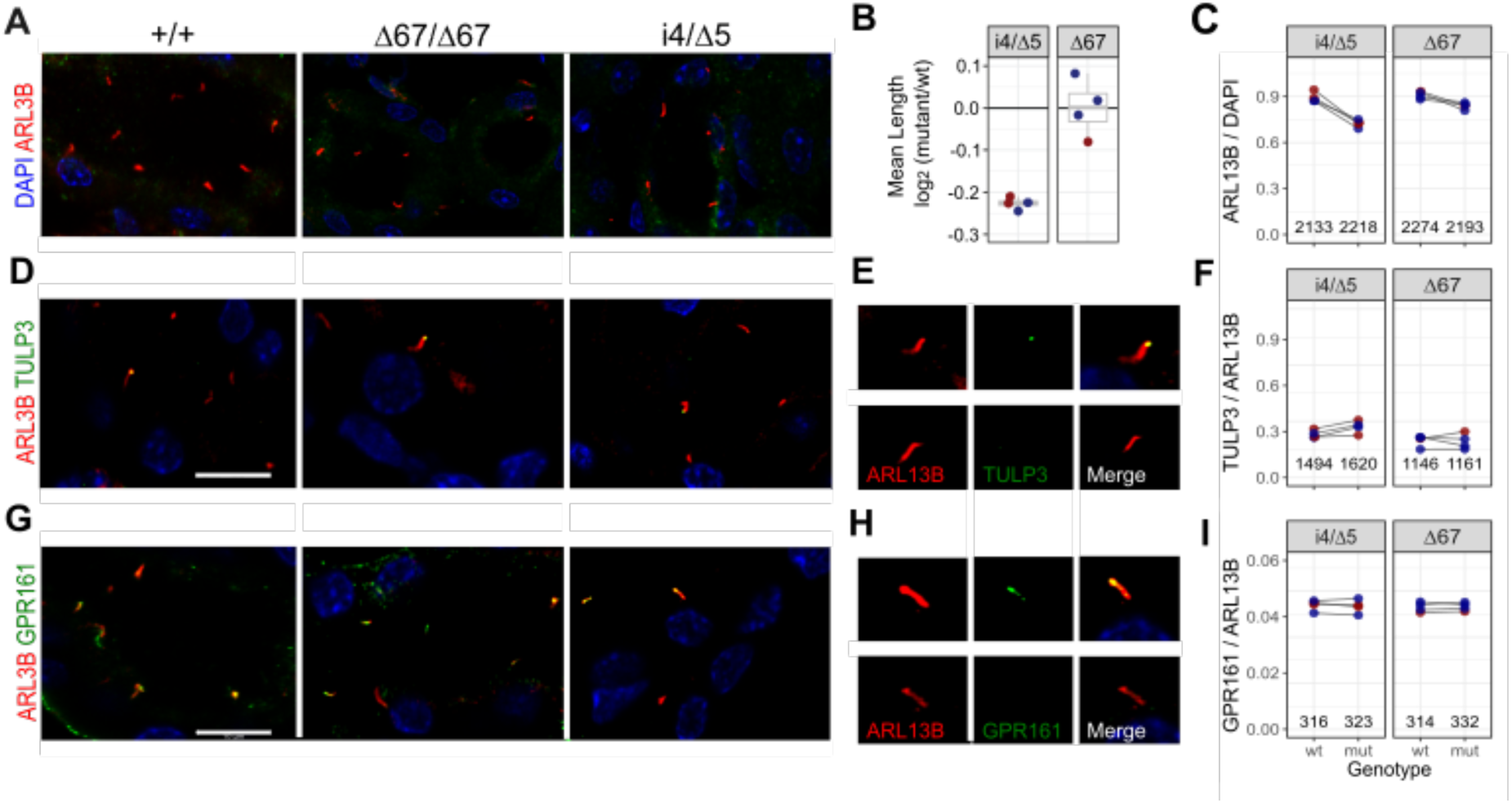
Hypomorphic *Tulp3* mutations affect frequency and length distribution of primary cilia. (**A**) Representative images of ARL13B immunofluorescence in primary cilia (red) with DAPI-stained nuclei (blue) from kidney sections of wild-type (+/+), i4/Δ5, and Δ67 mutant mice at postnatal day 20. Scale bar, 10 µm. (**B**) Mean cilium lengths for four littermate pairs, expressed as the log₂ ratio(mutant/wild-type). Female pairs red, males blue. (**C**) Ciliation frequency quantified as the percentage of ARL13B-positive cilia from 500-600 nuclei per sample. Lines connect control (left) and *Tulp3*-mutant (right) littermate samples. (**D**) Representative images showing localization of ARL13B (red) and TULP3 (green) relative to DAPI-stained nuclei (blue). Scale bar, 10 µm. (**E**) Magnified view of a single cilium showing single channel ARL13B (red), TULP3 (green), and merged channels to show localization of TULP3 to ciliary tip (top) or not (bottom). (**F**) Low but similar frequencies of TULP3 co-localization in ARL13B-defined cilia across genotypes. Lines connect littermate pairs. (**G**) Representative images showing ARL13B (red), GPR161 (green), and DAPI-stained nuclei (blue). Scale bar, 10 µm. (**H**) Magnified view of a single cilium showing single channel ARL13B (red), GPR161 (green), and merged channels to illustrate ciliary localization (top) or absence (bottom) of GPR161. (**I**) Quantification of GPR161 localization to cilia, expressed as the percentage of GPR161-positive cilia with ARL13B signal. Lines connect littermate pairs.

We next examined whether cilium localization of TULP3 or its cargo protein GPR161 were affected. TULP3 was infrequently localized to primary cilia in situ and neither decreased expression in i4/Δ5 (OR 1.2, p = 0.68, Fisher’s exact test) nor deletion of exon 4 (OR 0.98, p= 1.0) had a significant impact on TULP3 localization frequency (Figure 6D-F). GPR161 was more frequently detected (Figure 6G-I) but its localization to cilia was not significantly different in either i4d5 (OR = 0.99, p = 0.95, Fisher’s exact test) or Δ67 mutants (OR = 0.99, p = 1.0).

Together, these results suggest that while ciliogenesis is impaired, residual TULP3 is properly localized and able to support GPR161 trafficking and confirm that quantitative defects in ciliogenesis precede cyst formation in *Tulp3* mutant kidneys.

### *Tulp3* reduction does not suppress major *Zfp423* brain phenotypes

We previously showed that *TULP3/Tulp3* is a repressive target of the ZNF423 (mouse ZFP423) transcription factor in both mouse granule cell progenitors and a human medulloblastoma (DAOY) cell line and that simultaneous knockdown of both genes reversed a SMO translocation defect of *ZNF423* knockdown in DAOY cells [14]. To test whether germline mutations that decrease *Tulp3* level suppress *Zfp423* developmental phenotypes at an anatomical level, we crossed *Zfp423* null as well as two ZFP423 domain-specific mutations to the *Tulp3-*i4/Δ5 combination, in a coisogenic FVB/NJ background and measured weight and eight neuroanatomical parameters that we previously used to quantify a *Zfp423* allelic series [26] (Figure 7).

**Figure 7.**
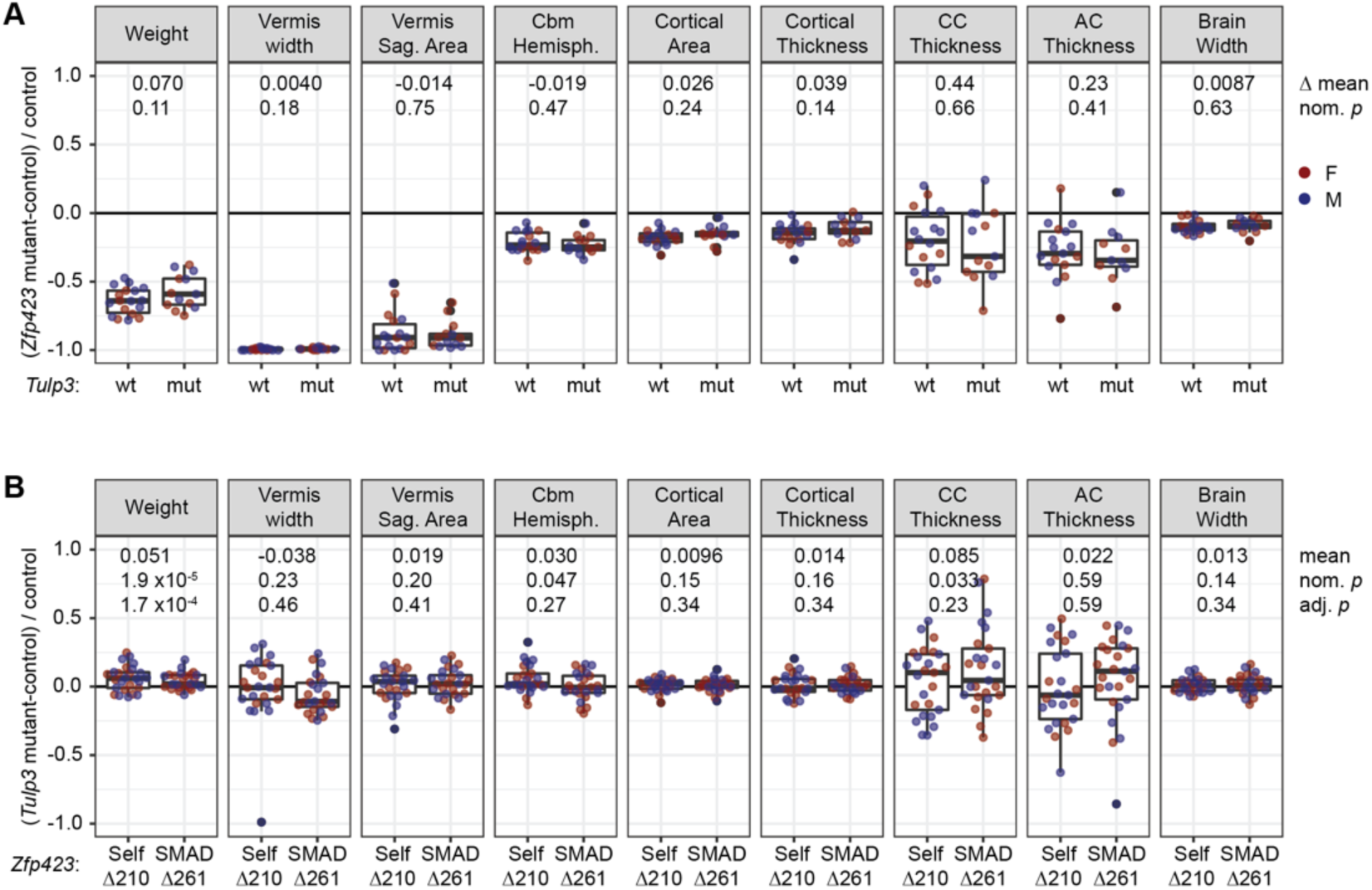
Genetic reduction of *Tulp3* does not suppress *Zfp423* null or hypomorphic phenotypes. (**A**) Major *Zfp423-*null phenotypes are not substantially improved by reduced *Tulp3* expression. Each point represents a *Zfp423* mutant-wildtype same-sex littermate pair, from crosses with *Zfp423^N507Tfs*43^* heterozygous and either *Tulp3* wild-type (wt, n=18), or *Tulp3* i4/Δ5 (mut, n=13) parents. Differences in the means between *Zfp423* and double mutants along with nominal *p*-values from Welch Two Sample t-test are given. (**B**) *Tulp3* reduction had only nominal effects on *Zfp423* hypomorph measures other than weight. Homozygotes for viable in-frame deletions of a self-association domain (Δ210) or SMAD-binding zinc fingers (Δ261) with viable reductions in *Tulp3* (i4/Δ5 or i4/i4) were compared to same-sex littermates homozygous for the same *Zfp423* mutation but wild-type for *Tulp3*. Self-association domain n=26 pairs, SMAD-binding domain n=25 pairs. Paired sample t-tests or Wilcoxon signed rank test (vermis width) before and after adjustment for family-wise error rate (Hommel’s method) are shown for the combined sample (n=51). Effects on weight were independently significant in each group.

We first compared *Zfp423*-null mutants (N515Tfs*43, also known as N507Tfs*43 or *Zfp423^em5Haml^*) with or without strongly reduced *Tulp3* expression (Figure 7A and S11_Table). Because *Zfp423* null homozygotes have reduced viability, we compared sex-matched mutant-wildtype pairs between two crosses of *Zfp423* heterozygotes, one in which all animals were *Tulp3* wild-type and one in which parents were *Tulp3*-i4/Δ5, resulting in offspring that were either i4/Δ5 or i4/i4. In the latter cross, paired samples had the same *Tulp3* genotype. We then compared the means of pair-wise differences between the two crosses. While seven out of nine measures trended toward improvement in the double mutant, neither individual measures nor the overall trend met even nominal significance at conventional statistical thresholds. While these results did not rule out a minor effect of *Tulp3* on *Zfp423* null phenotypes, nor an effect on some cell-level phenotype, they did not support the hypothesized genetic suppressor activity at the level of *Zfp423* brain structural abnormalities.

We also examined the effect of *Tulp3* on two viable and fertile alleles of *Zfp423* (Figure 7B and S12_Table). *Zfp423* encodes 30 C2H2 zinc finger domains and an additional putative non-canonical 4-cysteine zinc finger [28], clusters of which mediate several known physical interactions [22, 29–32], including a C-terminal homophilic interaction domain and SMAD protein-interacting region in zinc fingers 14-19. Both a 210-bp in-frame deletion (Δ210) that eliminated the 4-cysteine zinc finger region in the homophilic interaction domain and a 261-bp deletion (Δ261) that fused the first half of zinc fingers 15 to the second half of zinc finger 18 in the SMAD-interacting region resulted in quantifiable defects in weight and brain measures [26]. We used crosses in which both parents were homozygous for one of these two *Zfp423* domain deletions and heterozygous for *Tulp3*-i4 in one parent and *Tulp3*-Δ5 in the other to produce same-sex littermate pairs of *Zfp423* hypomorphs that are *Tulp3* wild-type or i4/Δ5. We observed nominal improvements in weight, anterior-posterior distance in cerebellar hemispheres, and thickness of the corpus callosum, but each effect had small magnitude and only weight survived correction for family-wise error rate across the nine measures. These results suggest that *Tulp3* reduction may have had a suppressing effect on some *Zfp423* anatomical phenotypes in hypomorphic variants, but that any effect sizes were quite small relative to the *Zfp423* main effects.

## DISCUSSION

Here we demonstrated a simple method for creating a quantitative allelic series in mice and identified distinct TULP3 levels required for viability, neuroanatomical measures, kidney homeostasis, and obesity and used that series to calibrate a qualitative allele affecting both expression level and protein composition. These results provide a generalizable strategy for efficient quantitative animal models of genetic disorders and identify new features of *Tulp3* pathology in mice.

From a single “one-pot” CRISPR/Cas9 editing experiment at exon 7, we recovered 6 functionally distinct alleles (counting only one of two presumptive nulls, Figure 1) targeting the polypyrimidine tract adjacent to an essential, frame-shifting exon. Null variants recapitulated expected embryonic lethal phenotypes, including polydactyly and exencephaly, while other recovered alleles that reduced *Tulp3* RNA and protein expression showed a range of phenotypes that correlated with reduction in expression level. Our approach should be extensible to other genes, though details may vary with the strength of cis-acting signals for exon inclusion. This approach allows more accurate modeling of pathogenic variants by calibrating the amount of genetic function required for measured phenotypes with much more precision than simple null and heterozygote collections in large-scale phenotyping centers.

Our results support different quantitative requirements for TULP3 among affected tissues. Measures across calibrated *Tulp3* genotypes allowed us to assess quantitative thresholds and dose-responses for several notable phenotypes (Figure 8). The data showed a low threshold (∼5% of control) to allow viability while avoiding polydactyly and severe neural tube defects, although the strongest viable genotype had measurable if slight increases in some morphological brain measures. Among viable genotypes, we found significant excess weight relative to littermates that expressed less than one-third of control TULP3 levels, with a slightly larger impact of exon 4 deletion relative to its measured expression level. Obesity is a cardinal feature of mutations in *Tub*, the close paralog of *Tulp3* and eponymous founding member of the gene family [33–36]. Whether and to what extent variants at either locus might modify phenotypic expression of the other remains to be tested. Notably, we found later onset and less severe kidney phenotypes for viable *Tulp3* variant combinations in our FVB-coisogenic models than previous reports from mice on C57BL/6 or mixed backgrounds, despite inclusion of very strong reductions in TULP3 expression level. These differences may include effects of genetic modifiers, but we note that presumptive null variants on FVB recapitulate prior phenotypes. This suggests that substitution variants with highly cystic kidneys by birth, such as K407I [37], represent severe loss of function. Further studies will be required to determine whether these differences indicate strain-dependent genetic modifiers or environmental differences. Our data also showed that kidney cystogenesis had a quantitative dose-response to TULP3 level across a range from ∼5-30% of control levels. Notably, this is a range that is not addressed in large consortia that assess knockout and heterozygote models. Prior to detectable cystogenesis, kidneys showed a decrease in the frequency of primary cilia and in length of observed cilia, at least in the strongest viable genotype, i4/Δ5, but we did not find a significant difference in the proportion of evident cilia that were stained for either TULP3 or GPR161. We also found no evidence for *Tulp3*-mediated suppression of *Zfp423* phenotypes predicted from cell culture assays. This highlights the importance of in situ validation of acute perturbation cell culture models of complex tissue-level phenotypes. These data provide a unique quantitative framework for modeling human *TULP3* effects in mice with a generalizable approach and for estimating quantitative effects of qualitative protein variants.

**Figure 8.**
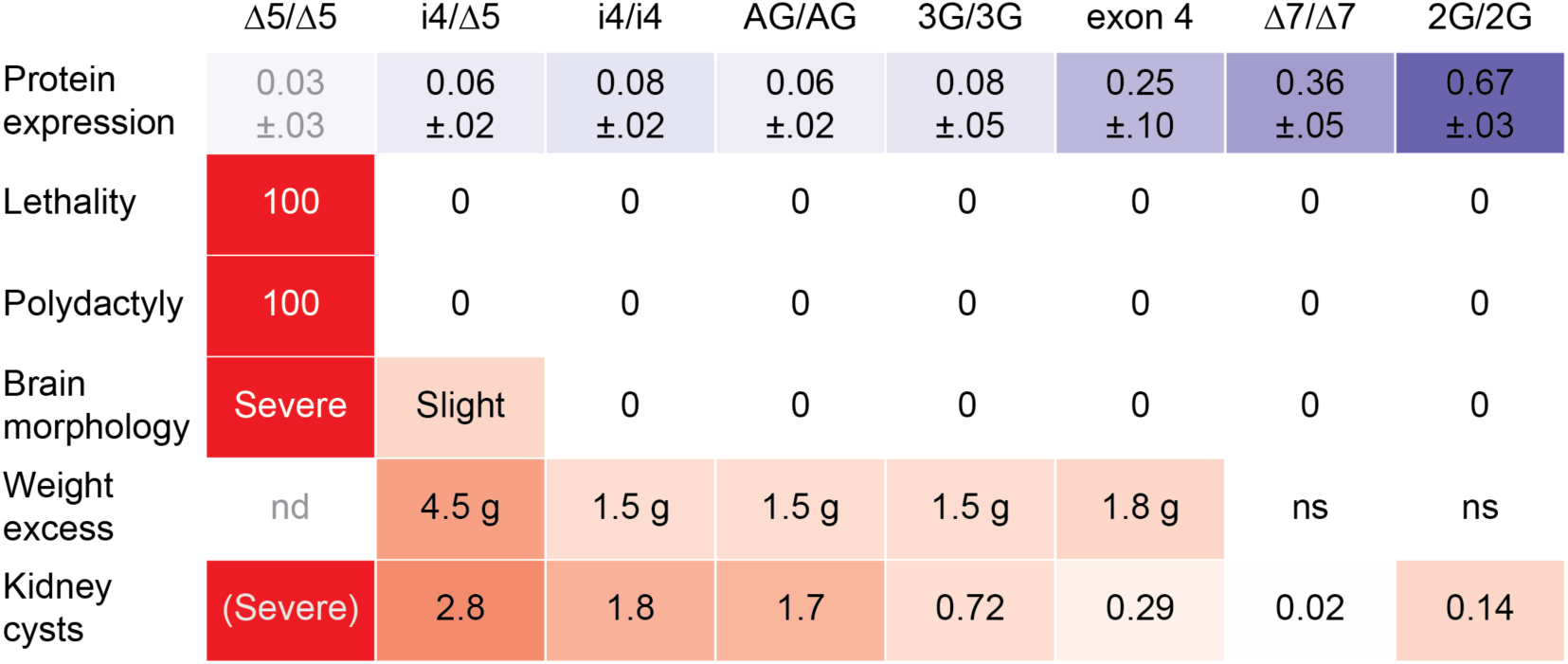
Summary of TULP3 expression-dependent effects. Heat map shows mean protein level in E14.5 embryo heads (top row, blue) inversely proportional to measured phenotypes (rows 2-6, red). Very low levels of TULP3 expression are sufficient to avoid lethality and polydactyly. Only the lowest viable expression level has residual brain morphological abnormalities relative to same-sex control littermates. Genotypes with less than ∼36% TULP3 expression show increased weight gain, with average values relative to same-sex littermates at ∼100 days shown. Cystic index in kidneys showed continuous graded response to TULP3; interpolated average at 200 days is shown; severity of the null inferred from embryonic and juvenile kidneys of earlier *Tulp3* models [16, 17]. nd, not determined; ns, no significant difference.

## MATERIALS AND METHODS

### Mice

FVB/NJ mice were purchased from The Jackson Laboratory. *Zfp423* mutant mice were previously reported [26] and maintained coisogenic on FVB/NJ. Mice were maintained in a specific pathogen free (SPF) facility on 12 h light, 12 h dark cycle in high-density racks with HEPA-filtered air and ad libitum access to water and food (LabDiet 5P06). All animal experiments were approved by the University of California San Diego Institutional Animal Care and Use Committee (UCSD-IACUC) under protocol S00219.

### Germline editing

Genome editing was contemporaneous with and essentially as described by Deshpande et al. [26]. Briefly, FVB/NJ one-cell embryos were injected with pre-assembled editing mixes in IDTE (10 mM Tris, 0.1 mM EDTA, pH 8.0) and transplanted into pseudo-pregnant females in the UC San Diego Moores Cancer Center Transgenic Mouse Shared Resource facility. Editing mixes were formulated by annealing tracrRNA and crRNA for ∼30 minutes in IDTE or 0.5x duplex buffer (0.5x is 15 mM HEPES, pH 7.5; 50 mM potassium acetate), incubating with diluted Cas9 for 5 minutes (exon 4) or HiFi Cas9 for 15 minutes (exon 7), diluting the resulting complex in IDTE, and supplementing with oligonucleotide pools (Ultramers) as donors for homology-dependent repair. Diluted editing mixes as injected contained 1.2 to 1.5 µM crRNA:tracrRNA, 30 ng/µl Cas9 or HiFi Cas9, and 2.4 to 3.2 µM pooled Ultramer oligonucleotides. Targeting sequences for each crRNA, with protospacer-adjacent motif (PAM, in parentheses), were: CAGCTGGACCGTCGATGCCT (AGG) and TAGGAGCACATCCCCGCAAC (CGG) for exon 4 and CCGGCTAGAAGAAATATCTG (AGG) for exon 7. Ultramer oligonucleotide sequences are given in Supplemental Table S1. All editing components were purchased from Integrated DNA Technologies (IDT). Tail biopsy DNAs from potential founder (G0) animals were used for PCR-based sequencing using primers in Supplemental Table S2. Edited lines were maintained coisogenic by serial backcross to FVB/NJ. Genotypes for stock maintenance and experimental subjects were determined by PCR assays (Supplemental Table S2). Nucleotide substitution allele (2G, 3G, and AG) used bidirectional-PCR amplification of specific alleles (biPASA) assay designs [38] unique to each allele.

### RT-PCR

RNA was extracted with Trizol reagent from E14.5 embryo heads and reverse transcribed with SuperScriptIII (Invitrogen). Exon inclusion assays used 35 cycles of PCR with primers in flanking exons (Supplemental Table S2) and resulting products resolved by gel electrophoresis.

### Western blots

Staged embryos were obtained by timed mating after 5 pm, with noon of the following day designated E0.5. Protein extracts from E14.5 heads were prepared in RIPA buffer (Teknova, R3792) supplemented with protease inhibitors (Sigma-Aldrich, P8340) from mutant and control littermate samples were separated in 13% acrylamide gels and electrophoretically transferred to nitrocellulose membranes (GVS, 1212590). After blocking in 5% non-fat dry milk (APEX, 20-241), membranes were exposed to rabbit anti-TULP3 ([2], gift of Dr. Jonathan Eggenschwiler) anti-phosphoprotein antibodies as a loading control (MilliporeSigma, P3300, P3430, P3555) followed by IR-680 conjugated donkey anti-mouse (LiCOR 926-68072) and IR-800 conjugated donkey anti-rabbit (LiCOR 926-32213) secondary antibodies. After washing 3x in TBST (20 mM Tris, 150 mM NaCl, 0.1% Triton X-100) retained antibodies were imaged on a LI-COR Odyssey imaging station and bands quantified using ImageStudio software (LiCOR).

### Anatomical measures

Sequential, paired samples were weighed ∼weekly for up 300 days. A small number of animals, predominantly females, developed circling behavior independent of genotype and were removed from the study. Simple brain morphological measures were made as described for *Zfp423* mutants [26]. Briefly, anesthetized mice were perfused with phosphate buffer followed by 4% paraformaldehyde. Brains were removed and post-fixed in 4% paraformaldehyde at 4°C overnight. External dorsal and block face coronal and sagittal images were captured on a Zeiss stereoscope with a Nikon Ds-Fi1 camera and measurements made in ImageJ software [39]. Kidneys were post-fixed in 4% paraformaldehyde overnight, then cryoprotected in 20% sucrose, 1X PBS, 0.5 mM EDTA. Tissues were embedded in O.C.T. compound (Fisher Scientific, 23-730-571) and flash frozen using 2-methylbutane cooled on dry ice. 15 µm sections were cut using a Leica CM3050S Cryostat at −20°C. After staining with hematoxylin and eosin, sections were photographed and cystic index was measured using CystAnalyzer software [27] using a 10 µm diameter setting.

### Cilium measures

Kidneys were prepared from sex-matched littermate pairs as above at postnatal day 20. 10 µm sections were cut at −20°C. Sections were washed in PBS three times for 5 minutes each and permeabilized with 0.1% Triton X-100 in PBS for 10 minutes. Sections were blocked with 5% donkey serum, 1% bovine serum albumin (BSA), and 0.1% Triton X-100 in PBS for 60 minutes at room temperature. Slides were incubated overnight at 4 °C with primary antibodies diluted in blocking buffer. Mouse anti-γ-tubulin (Sigma, T5326) and rabbit anti-ARL13B (Proteintech, 17711-1-AP) were used to measure cilia length. Mouse anti-ARL13B (Santa Cruz Biotechnology, sc-515784), rabbit anti-TULP3 (Proteintech, 13637-1-AP), and rabbit anti-GPR161 (Proteintech, 29328-1-AP) were used to assess the localization of TULP3 and GPR161. Sections were washed three times with PBS the following day and incubated with fluorophore-conjugated secondary antibodies (Invitrogen, A21203 and A21206) for 1 hour at room temperature in the dark. After three additional PBS washes, nuclei were stained with DAPI for 3 minutes, followed by a brief PBS rinse. Slides were mounted with Cytoseal 60 (Electron Microscopy Sciences, 18006), cover slipped and sealed. Fluorescence images were acquired at the UCSD Microscopy Core using a Leica SP8 confocal microscope equipped with Lightning Deconvolution. Image analysis was performed using LASX software on 5 µm Z-stack images.

### Statistical analyses

Data were visualized using ggplot2 with the ggbeeswarm package [40]. Data were assessed for normality using the Shapiro-Wilk test and evaluated for differences between same-sex littermates of opposite genotype using either parametric (t-test) or nonparametric (Wilcoxon signed rank test) for paired samples. Resulting nominal p-values were corrected for family-wise error rate (FWER) across phenotypes using Hommel’s method [25].

Survival analysis was performed using a Cox proportional hazards model in the survival package. All statistical tests were performed in R.

### Ethics statement

All animal experiments were approved by the University of California San Diego Institutional Animal Care and Use Committee (UCSD-IACUC) under protocol S00291.

## ACKNOWLEDGEMENTS

We thank Dr. Jonathan Eggenschwiler for the gift of TULP3 antibody. We thank Ella Kothari and Jun Zhao for mouse zygote injections performed in the UCSD Moores Cancer Center Transgenic Mouse Core Facility. We thank Oliver Zhang for genotyping assistance and reading the draft manuscript.This work was supported by grant R01 NS097534 from the National Institute of Neurological Disorders and Stroke to BAH. CAM was supported in part by a training grant from the National Cancer Institute, T32 CA067754. RZL was supported in part by an institutional grant from the National Institute of General Medical Sciences, R25 GM083275. The UC San Diego Transgenic Mouse Core is supported by NIH P30 CA023100 and P30 DK063491. The UC San Diego School of Medicine Microscopy Core is supported by P30 NS047101. The funders had no role in study design, data collection and analysis, decision to publish, or preparation of the manuscript.

## DATA AVAILABILITY

All graphical summary plots show individual measures as dots. Underlying tabular data are in supporting tables formatted as data frames.

---

**Supplemental S1_Figure.**
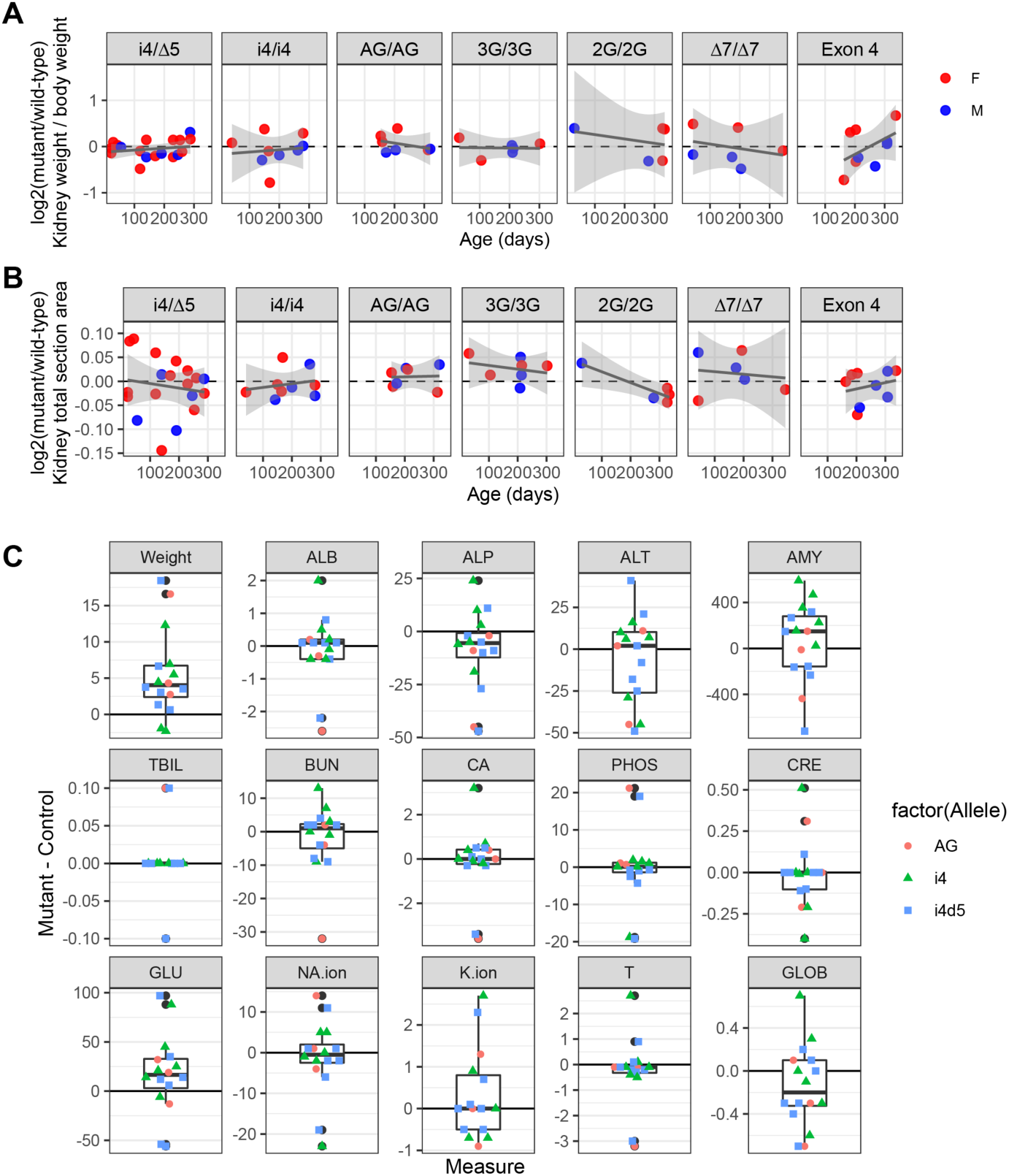
Additional paired sample kidney and blood chemistry measures. (**A**) Ratio of kidney weight to body weight was not significantly different between genotypes for any variant, p >0.2. (**B**) Total cross-sectional area of kidney was not significantly different between genotypes for any variant, p >0.2. (**C**) No striking differences in blood chemistries among 16 sex-matched littermate pairs. Standard blood chemistry panels from same-sex littermate pairs from strong *Tulp3* allele combinations found no consistent differences dependent on genotype, in contrast to their body weights, which confirmed the differences shown in Figure 3.

## Supporting Information captions

**S1_Table. Sequence of oligonucleotide donors used in editing exon 7.** Oligonucleotide donors used for genome editing in Figure 1C. The guide RNA sequences are given in the Materials and Methods.

**S2_Table. PCR primers used for screening and genotyping.** Left and right primers used to sequence potential founders and transmitted alleles for *Tulp3* exon 4 and exon 7 shown in Figure 1. The same exon 4 primers were used to genotype deletion alleles. Genotyping primers for biPASA assays included common “outer” primers with allele-specific inner primers for the reference (wt) and substituted (2G, 3G, or AG) allele. Short indel variants at exon 7 were typed using a 74-bp assay (Tx7.74). Exon 7 RT-PCR inclusion assay used a forward primer in exon 6 and reverse primer in exon 8.

**S3_Table. Normalized fluorescence values from Western blots.** Measures used in Figure 1H. Animal ID, genotype, and expression values from Western blots of extracts from embryonic day (E)14.5 samples. Mutant and heterozygote sample measures were adjusted to the intensity of their wild-type control littermate (pctWT).

**S4_Table. Normalized fluorescence values from Western blots.** Measures used in Figure 1I. Animal data and Western blot data for extracts from postnatal day 29 tissues. Mutant sample measures were adjusted to the intensity of their wild-type control littermate.

**S5_Table. Paired sample weight measures.** Longitudinal weight measures for paired littermates used to generate Figure 3. Logical variable (e.g., n75) describe whether a specific measure is the first past that postnatal day for that littermate pair. Removed indicates whether a measure was censored due to subsequent animal health problems for which weight change may have been a leading indicator independent of genotype.

**S6_Table. Paired sample brain measures.** Terminal weight and brain measures used in Figure 4. Measures follow those in Deshpande et al. [26]

**S7_Table. Paired sample kidney cystic index measures.** Animal characteristics and cystic index measures used in Figure 5.

**S8_Table. Paired sample blood chemistry measures.** Terminal weight and veterinary blood chemistry measures used in S1_Figure.

**S9_Table. Distribution of cilium lengths.** Length measures for 277 cilia across eight i4/Δ5 and Δ67 littermate pairs.

**S10_Table. Proportion of cilia with TULP3 and GPR161.** Proportions of ARL13B-stained cilia that had co-localized TULP3 or GPR161 immunofluorescence.

**S11_Table. Paired sample measures from *Zfp423*-null x *Tulp3*-i4/Δ5 cross.** Terminal weight and brain measures used in Figure 6A.

**S12_Table. Paired sample measures from *Zfp423*-hypomorph x *Tulp3*-i4/Δ5 cross.** Terminal weight and brain measures used in Figure 6B.

**S1_Figure. Paired sample kidney and blood chemistry measures.** (**A**) Ratio of kidney weight to body weight was not significantly different between genotypes for any variant, p >0.2. (**B**) Total cross-sectional area of kidney was not significantly different between genotypes for any variant, p >0.2. (**C**) Box-and-whiskers plots with individual data points for paired sample differences (mutant – littermate control) for Albumin (ALB), Alkaline Phosphatase (ALP), Alanine Transaminase (ALT), Amylase (AMY), total Bilirubin (TBIL), Blood Urea Nitrogen (BUN), Calcium (CA), Phosphorus (PHOS), Creatinine (CRE), Glucose (GLU), Sodium (NA-ion), Potassium (K-ion), Total Protein (TP), and calculated Globulin (GLOB).

